# SVF-derived extracellular vesicles carry characteristic miRNAs in lipedema

**DOI:** 10.1101/767483

**Authors:** Eleni Priglinger, Karin Strohmeier, Moritz Weigl, Carolin Lindner, Martin Barsch, Jaroslaw Jacak, Heinz Redl, Johannes Grillari, Matthias Sandhofer, Matthias Hackl, Susanne Wolbank

## Abstract

Lipedema is a chronic, progressive disease of adipose tissue with lack of consistent diagnostic criteria. The aim of this study was a thorough comparative characterization of extracellular microRNAs from the stromal vascular fraction (SVF) of healthy and lipedema adipose tissue. For this, we analyzed 187 extracellular microRNAs in concentrated conditioned media (cCM) and specifically in small extracellular vesicles (sEVs) enriched thereof by size exclusion chromatography. No significant difference in median particle size and concentration was observed between sEV fractions in healthy and lipedema. We found the majority of miRNAs located predominantly in cCM compared to sEV enriched fraction. Surprisingly, hierarchical clustering of the most variant miRNAs showed that only sEV miRNA profiles – but not cCM miRNAs – were impacted by lipedema. Seven sEV miRNAs (miR–16-5p, miR-29a-3p, miR-24-3p, miR-454-p, miR–144-5p, miR-130a-3p, let-7c-5p) were differently regulated in lipedema and healthy, whereas only one cCM miRNA (miR-188-5p) was significantly downregulated in lipedema. Comparing SVF from healthy and lipedema patients, we identified sEVs as the lipedema relevant miRNA fraction. This study contributes to identify the potential role of SVF secreted miRNAs in lipedema.

## Introduction

Lipedema is a chronic, progressive disease characterized by bilateral, symmetrical, disproportional deposition of adipose tissue in the extremities and buttocks^1^. Patients suffer from pain, reduced joint mobility, hematoma, edema and psychological impacts^2^. It was first described in 1940 as a connective tissue disorder, characterized by fluid being collected in the interstitium instead of entering into lymphatics^3^. This excess fluid in the interstitium potentially leads to growth of adipose tissue and hypoxia, which in turn might enhance angiogenesis of pathologic vessels^4,5^. The area of lymphatic vessels and the number of blood vessels were found increased in non-obese lipedema patients compared to controls^6^. Examination of adipose tissue from lipedema patients demonstrated hypertrophic adipocytes, crown-like structures and increased number of macrophages^6–8^.

Besides functioning as an energy storage, white adipose tissue (WAT) responds differentially to physiological and pathological metabolic changes by secreting a large diversity of proteins, hormones, lipids, non-coding RNAs – including microRNAs (miRNAs) – and extracellular vesicles (EVs)^9,10^. Small EVs (sEVs) are a fraction of 70-150 nm sized, membrane-enclosed particles, which contain cell-type specific proteins, enzymes, growth factors, cytokines, lipids, as well as coding and non-coding RNAs. It has been repeatedly reported, that WAT-derived vesicular miRNAs are involved in metabolic regulations^11,12^ and adipose tissue is considered a significant source of circulating sEV-miRNAs^11^. By acting in an autocrine, paracrine as well as systemic manner, these factors can contribute to metabolic abnormalities, modulation of osteogenic differentiation, inhibition of adipogenesis, adipocyte hypertrophy and infiltration of immune cells^13–15^. Many molecules released by adipose tissue originate from non-adipocyte cells, such as endothelial and immune cells present in the stromal vascular fraction (SVF) of adipose tissue^9,13^.

Critical issues are the unknown etiology of the disease and the lack of consistent diagnostic criteria leading to misdiagnosis in many cases. Clinicians must consider multiple criteria during the physical examination and conducting the medical history. A frequent comorbidity with 21.5% is cardiac disease^16^, however the risk of diabetes, dyslipidemia and hypertension is low despite an obese median BMI^1,17^.

To our knowledge, there are no blood-based parameters to identify lipedema patients so far. Due to this lack of systemic differences, we previously focused on the diseased tissue itself, specifically on the cellular components in which the disease is manifested and analyzed stromal vascular fraction (SVF) cells isolated from liposuction material of lipedema patients. The isolated cells showed a reduced adipogenic differentiation potential as compared to healthy controls, similar to Bauer et al.^18^. However, we observed a higher cell yield after isolation, which might be due to the enhanced cell number of mesenchymal (CD90) and the endothelial (CD146) positive cells^7^. Moreover, the proliferation capacity of adipose-derived stem/progenitor/stromal cells (Ki67+CD34+cells) was enhanced in lipedema tissue^8^.

MicroRNAs (miRNAs) are small non-coding RNAs that regulate gene expression through RNA interference. More than 70% of all human coding genes are under the control of microRNAs. Intracellular miRNA transcription and miRNA release from cells within EVs and protein complexes, which protect the RNA cargo from degradation, can inform about the onset and progression of human diseases^19^. Several studies have shown that either the overall number of EVs and/or the relative microRNA cargo can be altered in response to stress such as senescence or subtoxic liver damage^20,21^. Nevertheless, several studies suggest that the majority of extracellular microRNAs are contained in protein complexes rather than exosomes/microvesicles^22,23^.

Based on our previous knowledge about the relevance of SVF cells in lipedema and the potential link between extracellular microRNA profiles and disease phenotypes, the aim of this study was to perform a thorough characterization of the extracellular microRNA fraction produced by SVF in healthy (control) individuals and lipedema patients. For this, we analyzed 187 extracellular microRNAs in the concentrated conditioned media (cCM) and specifically in sEVs isolated thereof by size exclusion chromatography, to determine the relevant fraction containing the lipedema discriminating microRNAs.

## Methods

### SVF isolation

The collection of human adipose tissue was approved by the local ethical board (application/approval number 200, 12.05.2005 and 19.05.2014), and written patient’s consent. Table 1 provides patient characteristics. Subcutaneous adipose tissue was obtained during routine outpatient liposuction procedures from the hips and outer thighs (“saddlebags”) under local tumescence anaesthesia. Tumescence solution (per liter) contained 3.3 mg Volon-A (Dermapharm), 1 vial Suprarenin 1 mg/mL (Sanofi-Aventis), 15 mL bicarbonate 8.4% (Fresenius Kabi) and 23.3 mL Xylocaine 1% (Gebro Pharma GmbH). The harvesting cannulas were triport and 4 mm in diameter (MicroAire System power-assisted liposuction). 100 mL liposuction material was transferred to a blood bag (Macopharma, Langen, Germany) and washed with an equal volume of phosphate buffered saline (PBS) to remove blood and tumescence solution. Next, for tissue digestion PBS was replaced with 0.2 U/mL collagenase NB4 (Nordmark, Uetersen, Germany) dissolved in 100 mL PBS containing Ca^2+^/Mg^2+^ and 25 mM N-(2-hydroxyethyl)piperazine-N′-(2-ethanesulfonic acid) (HEPES; Sigma, Vienna, Austria), resulting in a final collagenase concentration of 0.1 U/mL. The blood bag was incubated at 37 °C under moderate shaking (180 rpm) for 1 h. The digested tissue was transferred into 50 mL-tubes (Greiner, Kremsmünster, Austria). After centrifugation at 1200 x*g* for 7 min the supernatant was removed and the cell pellet was incubated with 100 mL erythrocyte lysis buffer for 5 min at 37 °C to eliminate red blood cells. The supernatant after centrifugation for 5 min at 500 x*g* was aspirated and the cell pellet was washed with PBS and filtrated through a 100-μm cell strainer (Greiner). After another centrifugation step at 500 x*g* for 5 min the supernatant was removed. The isolated SVF was resuspended in medium filtrated (0.22 μm; Merck, Vienna, Austria) before: DMEM-low glucose (Lonza, Vienna, Austria) containing 10% fetal calf serum (FCS; Sigma, Vienna, Austria), 2 mM L-glutamine (Lonza). Cell number was determined using trypan blue exclusion and quantification in a cell counter (TC-20, Bio-Rad, Vienna, Austria).

**Table 1:**
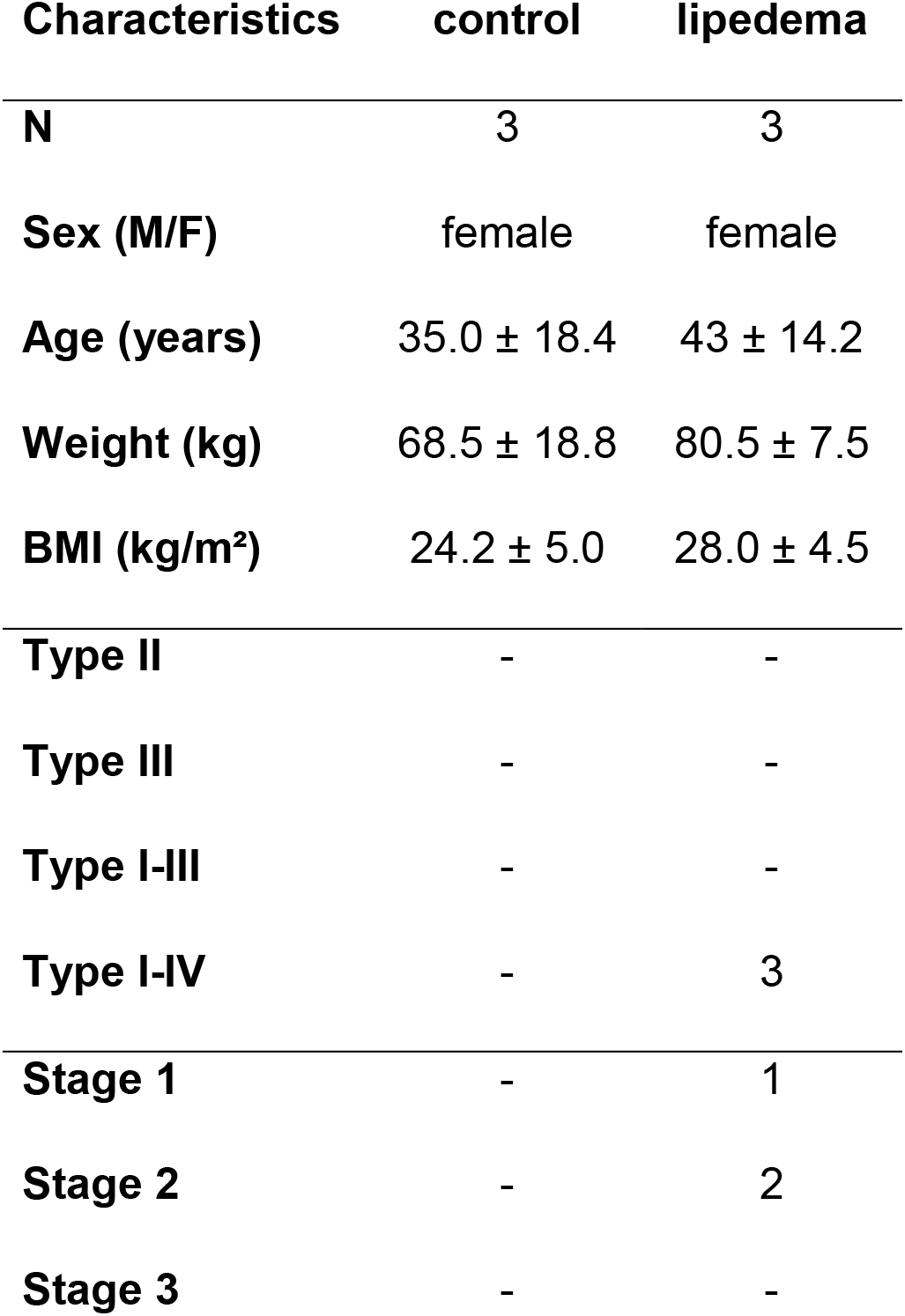
Characteristics of control and lipedema patients. The cohort underwent liposuction in this study were characterized regarding sex, age, weight and BMI and for lipedema patients the type and stage of lipedema. Data are presented as mean±SD.

### Collection of cell supernatants

For miRNA analysis, DMEM-low glucose containing 10% FCS, 2 mM L-glutamine was filtrated using 0.22 μm filter cups to remove large EVs. 6×10^6^ freshly isolated SVF cells were seeded in 22 mL filtrated medium in a T175 flask. The conditioned medium (CM) was collected after 24 h in a 50 mL-tube and centrifuged at 500 x*g* for 15 min (Eppendorf, 5804R; Vienna, Austria) to remove cellular debris. After transfer to a new tube (VWR, Vienna, Austria) and centrifugation at 14 000 x*g* for 15 min at 4°C (Heraeus Multifuge X3R; Thermo Fisher Scientific, Vienna, Austria) the supernatant was filtrated through a 0.8 μm strainer (VWR) and immediately frozen and stored at −80°C until further analysis.

### Processing of supernatants for microRNA analysis

CM was concentrated to 1 mL using centrifugation-based ultrafiltration (cCM). Briefly, 20 mL of CM were transferred to an Amicon 30 kDa ultrafiltration column and centrifuged at room temperature for 15 minutes. Residual volumes were measured and diluted to 1 mL using sterile PBS solution. Small extracellular vesicles (sEVs) were subsequently enriched from cCM by size exclusion chromatography using qEV70s single columns (Izon): 150 μL of cCM were loaded based on recommendations from the manufacturer, and fraction 8 – 11 were collected and pooled to obtain the fraction of enriched sEVs. Nanoparticle tracking analysis (NTA) was performed on all sEV samples to determine particle size and concentration.

### Nanoparticle tracking analysis (NTA)

For determination of size and concentration of particles CM, cCM and sEV fractions obtained by SEC were analyzed on Zetaview PMX 110 V3.0 particle analyser as previously described^24^ (Particle Metrix GmbH, Meerbusch, Germany).

### Total RNA extraction

Total RNA extraction was performed using 200 μl cCM or 200 μL pooled sEV fractions together with the miRNeasy mini kit (Qiagen). Synthetic oligonucleotides obtained from the miRCURY Spike-In kit (Qiagen) were added to the Qiazol lysis buffer before homogenization of the cCM/sEV samples. Glycogen (5 mg/mL) was added to the chloroform extract at 1:100 dilution to enhance precipitation. All other steps were performed according to the recommendations of the manufacturer. Total RNA was eluted in 30 μL nuclease-free water and stored at −80°C in low-bind tubes (Eppendorf) until further analysis.

### MicroRNA analysis

Reverse transcription (RT) of total RNA to cDNA was performed in 50 μL reaction volumes using the miRCURY RT kit (Qiagen) and 10 μL total RNA as input. The RT mix was incubated at 42°C for 60 min. The resulting cDNA was stored at −20°C until further analysis. qPCR reactions were prepared using the miRCURY SYBR® Green mastermix and a final cDNA dilution of 1:50. PCR amplification was performed in 384-well pick&mix plates (Qiagen) with customized selection of 192 LNA-enhance primers to detect 187 endogenous microRNAs and 5 spike-in controls per sample. The mastermix/cDNA sample was added to 384-well plates using an epMotion liquid handling robot (Eppendorf). Following the preparation of plates an incubation at 4°C for at least 60 min was performed before the plates were analyzed on a LightCycler 480 II (Roche) using 45 cycles and a temperature profile recommended by the manufacturer for miRCURY SYBR® Green mastermix. Cq-values were called using the second-derivate maximum method. Data quality was assessed using spike-in controls (Supplementary Fig. S2) to determine RNA extraction efficiency, enzymatic inhibition, and overall variability. Data normalization was performed using the global mean (i.e. average Cq-value for all endogenous microRNAs with Cq<35) to obtain delta Cq-values.

### Statistical analysis

Exploratory data analysis was performed using ClustVis. Global-mean normalized delta Cq-values (dCqs) were used for clustering analysis based Euclidean distance and complete linkage. Unit variance row scaling was applied. Statistical analysis was performed using unpaired two-sided t-tests.

## Results

### Clinical characteristics and sample collection

The cohorts that underwent liposuction in this study were all female, 3 healthy (=control) and 3 lipedema (=lipedema) patients. Average age was 35.0 ± 18.4 for the control and 43 ± 14.2 years for the lipedema group. The weight and body mass index of the control group was 68.5 ± 18.8 kg and 24.2 ± 5.0 kg/m^2^ and 80.5 ± 7.5 kg and 28.0 ± 4.5 kg/m^2^ for the lipedema group. The three lipedema patients were categorized as type I-IV and stage 1 or 2 (according to Dr. Barsch and Dr. Sandhofer; Table 1). For collection of the samples (cCM and sEVs) all steps from liposuction (site, anesthesia, cannula) to SVF isolation, cultivation and preparation were performed consistently for each sample (Fig. 1).

**Figure 1.**
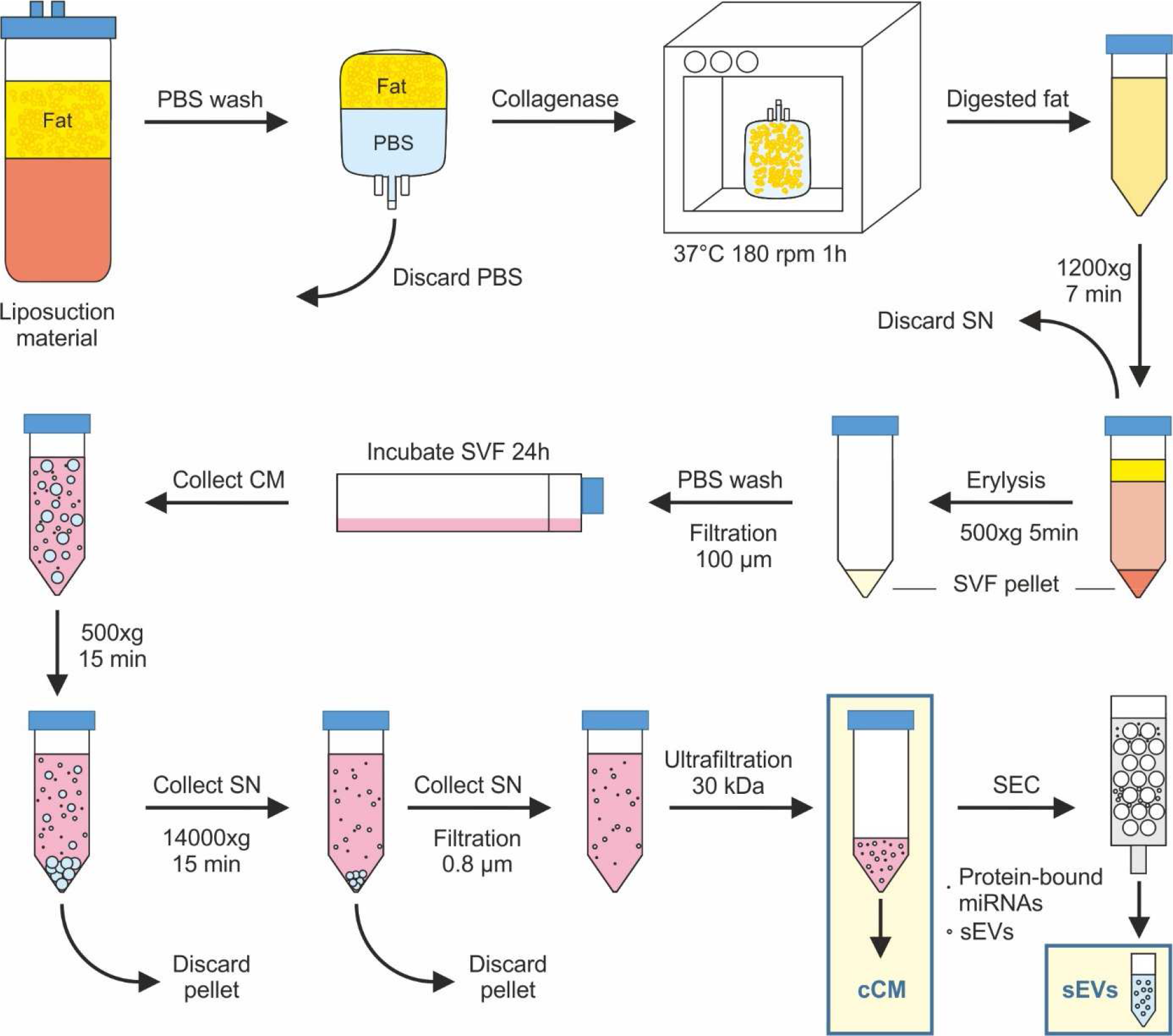
Experimental setup for SVF isolation, collection of cell supernatants and processing of supernatants for miRNA analysis. Subcutaneous adipose tissue was obtained during routine outpatient liposuction procedures. The liposuction material was washed and digested with collagenase. After centrifugation of the digested fat, the cell pellet was incubated with erythrocyte lysis buffer. After another centrifugation step, the cell pellet was washed and filtered and the isolated stromal vascular fraction (SVF) was incubated for 24h in sterile filtered media. Conditioned media (CM) was collected and after two centrifugation steps and filtration, it was concentrated using centrifugation-based ultrafiltration to obtain concentrated CM (cCM). Small extracellular vesicles (sEVs) were enriched from cCM by size exclusion chromatography (SEC).

### sEVs do not change in size or number in lipedema patients

In order to determine the relevance of SVF miRNAs in lipedema, we collected CM of adipose tissue derived SVF cells. NTA was employed to analyze particle size and concentration. By ultrafiltration a 17-fold enrichment of particles was achieved in the hereinafter called cCM (Supplementary Fig. S1 a). sEVs were successfully separated from proteins of the cCM by SEC (Supplementary Fig S1 b). Fractions 8-11 showed the highest particle count and were pooled for further analysis. The particle diameter did not differ between cCM and sEVs (Supplementary Fig S1 c and d), indicating a comprehensive sEV enrichment. Particle size obtained by size exclusion chromatography ranged between 50 nm and 700 nm. The median particle size was determined to be 165 nm (+/− 2.85) for the control group and 167 nm (+/− 10.78) for the lipedema group (Table 2). Similarly, no significant difference in particle concentration was observed between both groups, despite a trend towards higher concentrations in the lipedema group (FC=1.41).

**Table 2.** Particle number and size of purified extracellular vesicles (EVs) from control and lipedema analyzed by nanoparticle tracking analysis. Nanoparticle tracking analysis was applied to the small extracellular vesicles (sEVs) samples from controls and lipedema patients to determine concentration and size of purified nanoparticles. No significant differences were observed in particle concentration and particle size between controls and lipedema patients.

| Group |  | Particle number<br>[Particles/ul] | Median particle<br>size [nm] | Mean particle<br>size [nm] |
| --- | --- | --- | --- | --- |
| control 1 |  | 1.63E+14 | 161 | 182 |
| control 2 |  | 2.87E+14 | 166 | 186 |
| control 3 |  | 2.73E+14 | 168 | 191 |
| lipedema 1 |  | 5.07E+14 | 153 | 167 |
| lipedema 2 |  | 2.97E+14 | 179 | 197 |
| lipedema 3 |  | 2.20E+09 | 168 | 187 |
| control vs. lipedema | Effect Size<br>(fold change) | 1.11 | 1.01 | 0.99 |
| control vs. lipedema | p-value<br>(2-sided t test) | 0.87 | 0.85 | 0.81 |

### The majority of microRNAs are more abundant in cCM compared to sEV fraction

192-plex RT-qPCR panels were used to quantify 187 microRNAs and 5 control assays in all 12 samples (6x cCM and 6x sEV). Spike-in controls showed homogenous values across all 12 samples with low variability, demonstrating low analytical variability, no enzymatic inhibition and overall high quality of the analysis (Supplementary Fig. 2). In the cCM fractions 182/187 were above the limit of detection, while 133/187 miRNAs were detectable in the sEV fractions (and cCM). This set of 133 miRNAs was used for all further statistical analyses. The global mean normalized miRNA levels in cCM samples were interpreted as the total miRNA signal, since the analysis captures miRNA cargo in EVs as well as protein complexes. In order to determine how much of the total signal came from the sEV fraction, the sEV enriched miRNA signals were related to the cCM miRNA signals and expressed as percentage of microRNA signal derived from sEV. Fig. 2 shows the top and bottom 20 miRNAs (Fig. 2a and b) in sEV and cCM. Supplementary Table 1 presents the results for all 133 miRNAs. We identified three miRNAs, for which more than 90% of the signal in cCM originated from the sEV enriched fraction (miR-144-3p, miR-144-5p, miR-190a-5p), and 6 further miRNAs where >50% of the signal was derived from the sEV fraction. However, 83 out of 133 miRNAs were highly enriched in the non-sEV fraction as <10% of the total signal was obtained from the sEV fraction. This set of miRNAs included for example the miR-30 family and miR-23-24-27 family of miRNAs.

**Figure 2.**
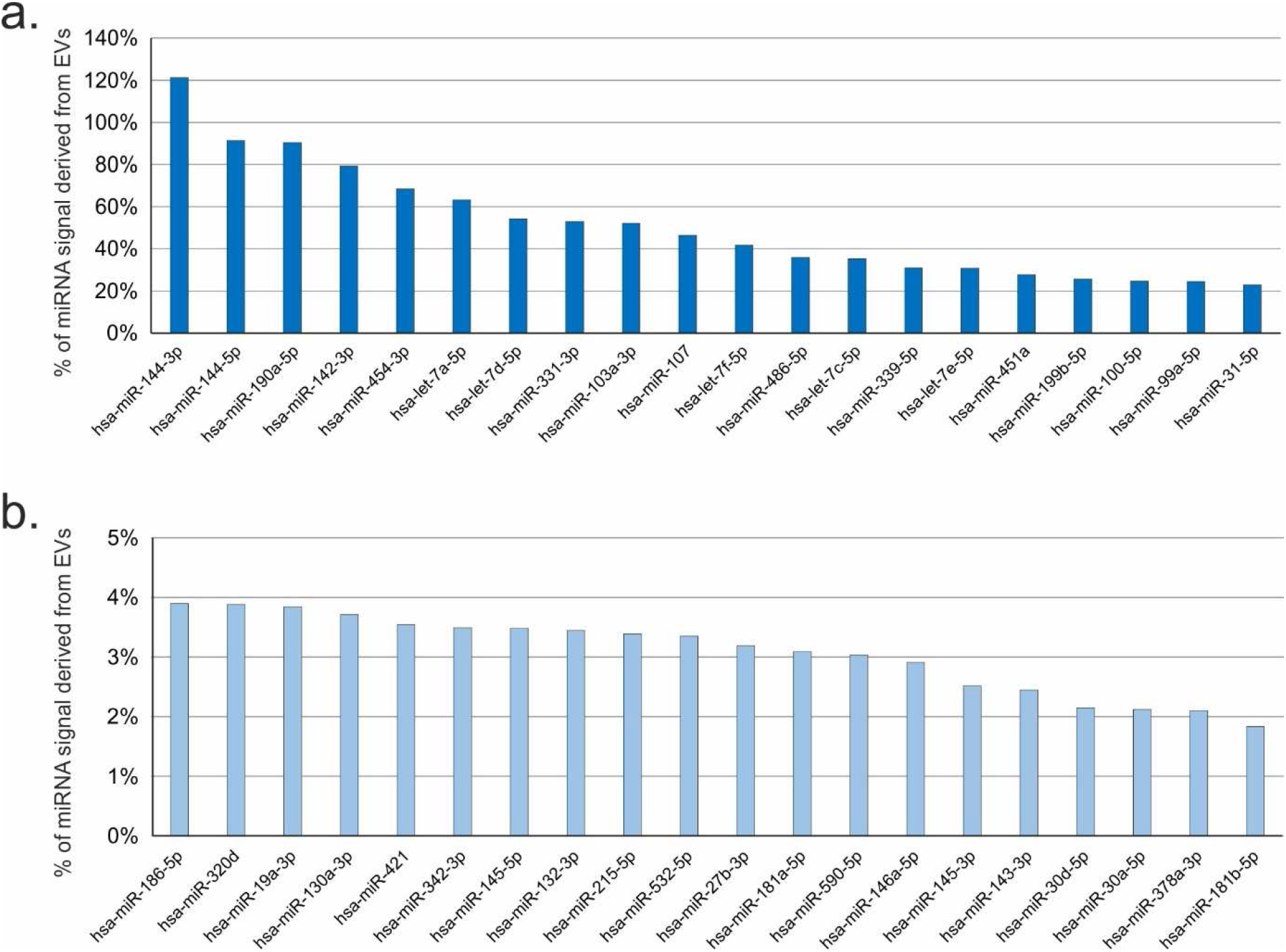
Analysis of microRNA signal origin. microRNAs signals obtained in small extracellular vesicles (sEVs) were compared against the total microRNA signal in concentrated conditioned media (cCM) and expressed as %. a) Top 20 miRNAs with highest EV-derived signal. b) Bottom 20 miRNAs with signals mostly derived from non-EV miRNAs.

### Clustering and differential expression analysis based on cCM and sEV microRNA profiles

The most variant miRNAs in the cCM and sEV enriched fraction were selected based on the coefficient of variation and used for hierarchical clustering (Fig. 3). While the variability in the cCM miRNA data was not primarily influenced by the disease (Fig. 3a, no clusters corresponding to the groups), this was the case for sEV enriched miRNA data, where 2 control samples clustered, and 2 lipedema samples clustered. Next, we performed statistical analysis to identify potentially regulated miRNAs. In general, the overall trend in up- and down-regulated miRNAs between lipedema and control was balanced for both the cCM and sEV enriched fraction (Fig. 4a and 4b). When applying a cut-off of p<0.05, we observed one microRNA (miR-188-5p, p=0.047) to be significantly down-regulated in the cCM fraction of lipedema patients compared to controls (Fig. 5h). Interestingly, this miRNA was not detected in the sEV fraction. Vice-versa, 7 miRNAs (3 up: miR–144-5p, miR-130a-3p, let-7c-5p; 4 down; miR–16-5p, miR-29a-3p, miR-24-3p, miR-454-p,) were identified to be significantly regulated in the sEV enriched fraction (Fig. 5a-g).

**Figure 3.**
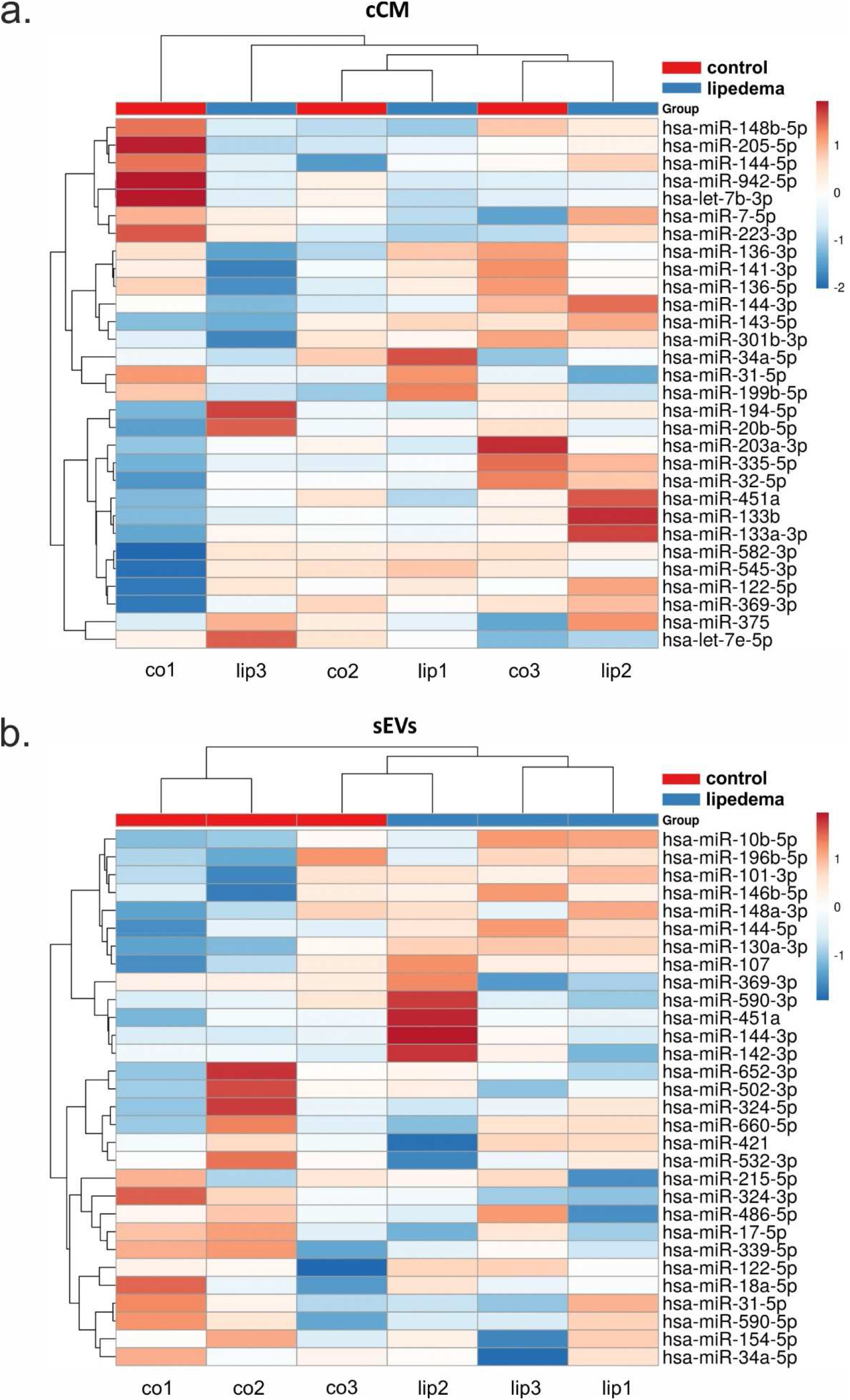
Hierarchical clustering of samples based on miRNA profiles observed in concentrated conditioned media (cCM) or small extracellular vesicles (sEVs) from control (co) and lipedema (lip). The 30 most variant (according to coefficient of variation) miRNAs in cCM (a) and sEVs (b) were used for clustering analysis (Pearson correlation, average linkage). Rows are centered and unit variance scaling is applied to rows. There were no clusters corresponding to the groups in the cCM miRNA data (a), while in the sEV enriched miRNA data 2 control samples clustered and 2 lipedema samples clustered (b).

**Figure 4.**
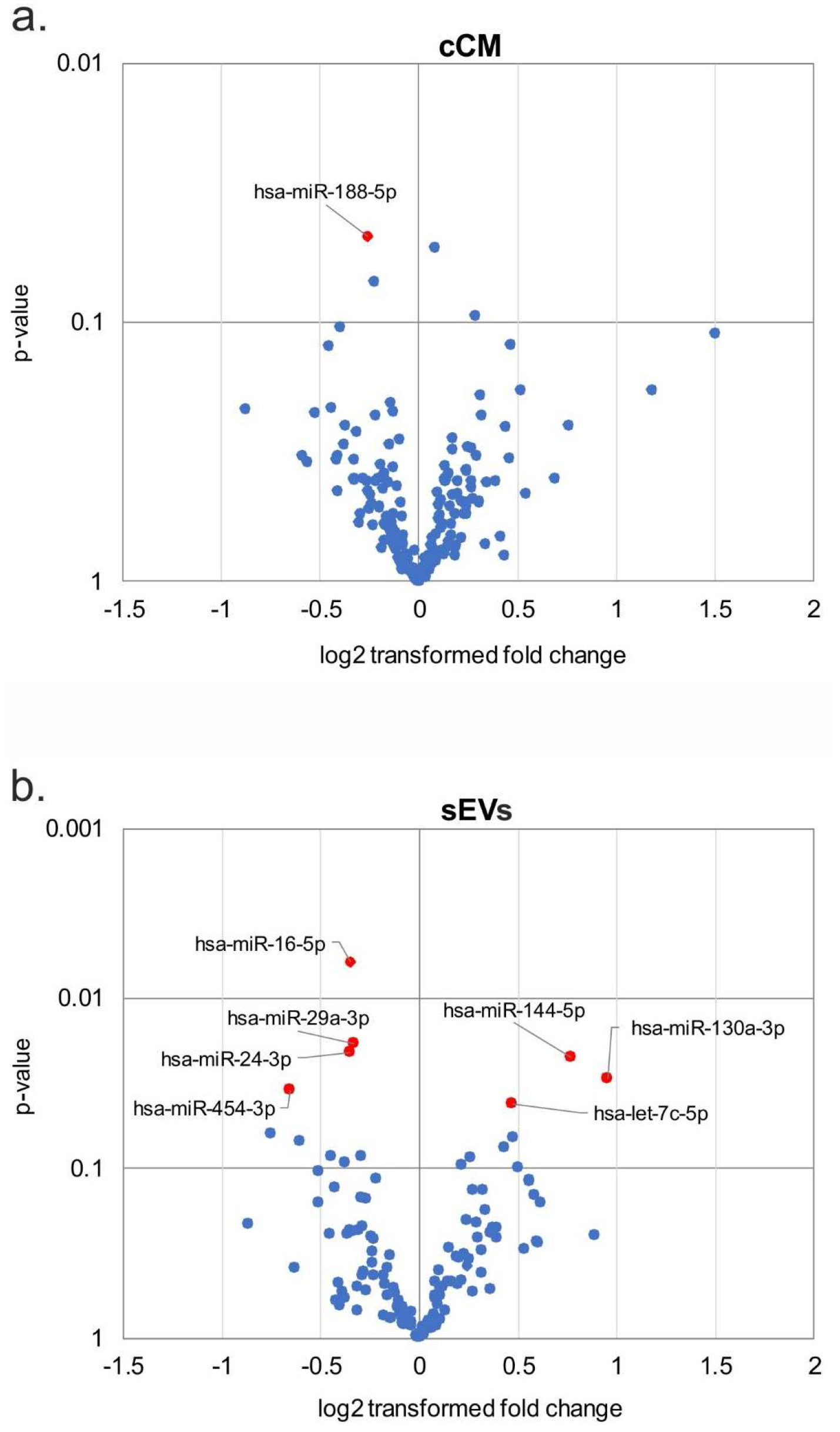
Differential secretion of extracellular miRNAs between control and lipedema. 182 microRNAs were analyzed by RT-qPCR in concentrated conditioned media (cCM) (a) and small extracellular vesicles sEV (b) fractions. Volcano plots depict the log2 transformed fold change and p-value for the measured miRNAs. The overall trend in up- and down-regulated miRNAs between lipedema and control was balanced for both the cCM (a) and sEV (b) fraction. P-value < 0.05 are highlighted in red. n=3 per group, unpaired two-tailed t-test.

**Figure 5.**
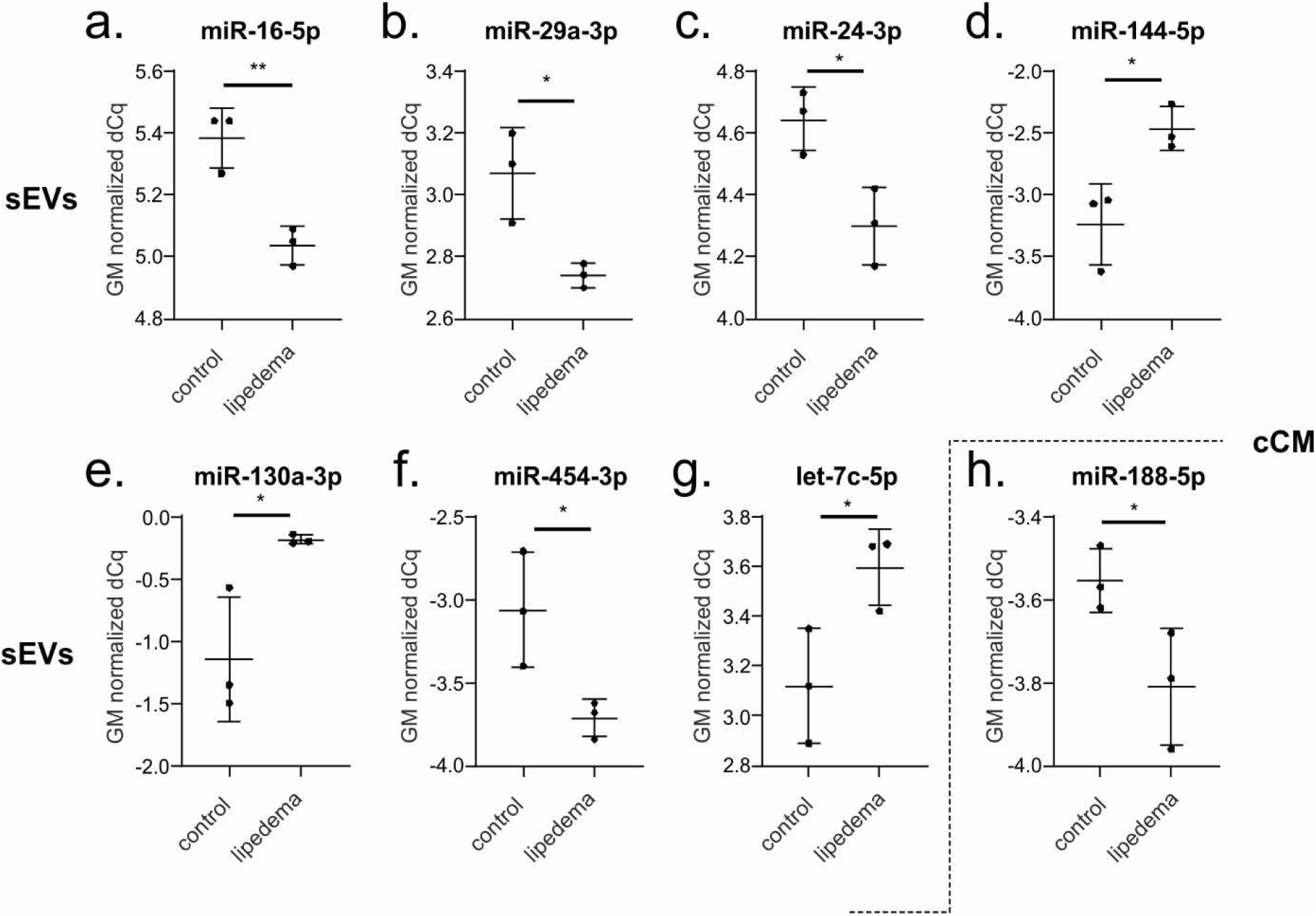
Scatterplots depicting global mean normalized levels of significantly regulated microRNAs in small extracellular vesicles (sEVs) and concentrated conditioned media (cCM) fraction. When applying a cut-off of p<0.05, 7 miRNAs (3 up, 4 down) were identified to be significantly regulated in the sEV fraction: miR-16-5p (a), miR-29a-3p (b), miR-24-3p (c) and miR-454-3p (f) were downregulated in lipedema patients compared to controls, miR-144-5p (d), miR-130a-3p (e) and let-7c-5p (g) upregulated. One miRNA was identified to be significantly downregulated in the cCM fraction of lipedema patients compared to controls (h). n=3 per group, unpaired two-tailed t-test, ** p<0.01, * p<0.05.

## Discussion

Since lipedema does not respond to lifestyle changes nor is there a treatment option other than liposuction of the diseased adipose tissue, a biomarker that would allow for identification of this disease and the prescription of liposuction by healthcare providers would be desirable. To date, diagnosis is based on clinical anamnesis, analytical weighing measurement and imaging of tissue composition by ultrasound. Additionally, non-invasive imaging such as MRI based quantification has been investigated. Adipose and sodium content in lipedema patients was found elevated in subcutaneous fat and skin, which hence has been suggested as potential biomarker for lipedema^25^. Since the disease is usually triggered by hormonal changes in females, a genetic susceptibility has been suggested^26^ (e.g. Williams-Beuern syndrome)^27^, however so far, no single gene causing the disease has been identified. The molecular load of EVs reflects the (patho-) physiological status of its original cells and thus can serve as a diagnostic tool, where sampling via liquid biopsies brings immense benefit to the physician and patient^28^. Besides the cell type-specific proteins, lipids and different classes of nucleic acids, EV contained miRNAs were found to be potent regulators in biological processes and in various diseases^29,30^. Numerous studies have already shown that the EV composition in diabetes, obesity, atherosclerosis, neurodegenerative diseases^31^, different cancers^32–34^ and renal damage^35–37^ distinguishes health and disease, which recommends them as potential diagnostic biomarkers. In fact, recent clinical trials have identified EVs as biomarker of prostate cancer^38^, Parkinson’s disease^39^ and difficult-to-treat arterial hypertension^40^.

Since miRNAs are crucial regulators and involved in the progression of human diseases, our present study focused on extracellular miRNA in lipedema. Previously we identified that SVF cells from diseased subcutaneous tissue are altered in terms of adipogenesis, cell content and cellular subtype composition^7^. Thus, we sought to analyze the miRNA secretory profile of this cell population in lipedema versus healthy subjects (control). Considering the recent discussion about the relevance of extra- and intravesicular miRNAs^41^, we differentially analyzed the total and EV-contained miRNAs secreted by SVF from lipedema patients and healthy controls. Turchinovich et al. postulated that extracellular miRNAs are predominantly non-vesicular, potentially a byproduct of dead cells but associated mostly with proteins^22^. In our study, the majority of the characterized miRNA were indeed located extra-vesicular. However, we found that only the EV-contained miRNA profile allowed discriminating healthy from lipedema, which was not the case for the secreted miRNA in the cCM, consisting mainly of extra-vesicular miRNAs. This demonstrates the importance of detailed characterization and especially the relevance of the EV cargo when characterizing the cellular status of a diseased tissue. We identified in our sEV preparations a significant reduction in miR–16-5p, miR-29a-3p, miR-24-3p, and miR-454-p, whereas miR–144-5p, miR-130a-3p and let-7c-5p were significantly increased in SVF derived sEVs in lipedema compared to healthy controls. In the cCM, we found a significant reduction of one single miRNA, miR-188-5p in lipedema. MiR-188-5p has been described as a marker for age-related switch from osteogenic to adipogenic differentiation^42^. Further, miR-188-5p may have the potential to regulate migration and support of vascularization by MSC, as previously described for a murine choroidal neovascularization model^43^. Deregulation of miR-188-5p might hence contribute to the endothelial barrier dysfunction and lymphangiopathy, which has been described in lipedema^44^. This suggests also a potential involvement of the mesenchymal stem cell populations of the SVF in the progression of lipedema, which are participating in the formation and stabilization of vessels^45^. It has been demonstrated that the total number of circulating particles (incl. their miRNA content) can be altered in disease^30,46^. In our study, particle number and size of enriched sEVs, analyzed by NTA, showed no difference between healthy and lipedema. The majority of single miRNAs showed higher abundance in cCM than in sEVs and only 9 of the analyzed miRNAs showed >50% signal coming from sEVs. miR-142-3p (80% sEV signal), was recently shown by another group to be selectively packaged in sEVs by a panel of oral squamous cell carcinoma cell lines, which resulted in lower intra-cellular levels and promoted malignant changes in these cells via increase of TGFBR1 expression^47^.

Several miRNAs have recently been found to regulate adipose tissue biology and play a role in the development of obesity and related metabolic complications^48–51^. Let-7 is one of the first miRNAs discovered. Knockout of let-7 family improved glucose tolerance in mice with diet-induced obesity^52^. In the present study, we observed a significant enhancement in let-7c-5p contents in sEVs from lipedema patients compared to healthy controls. Although we found significant increase in let-7c-5p, there is a consensus that lipedema does not favor the onset of diabetes, quite the opposite, lipedema negatively correlates with the development of insulin resistance. This is in line with observations by Pinnick et al. who found at cellular and miRNA level intrinsic differences between abdominal and gluteal adipose tissue. This results in an opposing metabolic disease risk, which is reduced for the gluteal region – the site affected in lipedema^53,54^. Let-7c has been associated with regulation of macrophage polarization^55^. After traumatic brain injury, let-7c-5p protected from neuroinflammation and attenuated activation of microglia/macrophages in a murine model of traumatic brain injury^56^. This reported impact on macrophage polarization is also in consistence with another study demonstrating that miR-9, miR-127, miR-155, and miR-125b induce M1 polarization, while miR-124, miR-223, miR-34a, let-7c, miR-132, miR-146a, and miR-125a-5p promote M2 polarization in macrophages^57^. Since lipedema is an inflammatory disease involving infiltration of macrophages forming crown-like structure around adipocytes^9^, the interplay of macrophages and local tissue cells of the diseased subcutaneous fat, especially the contribution of EV miRNA communication requires in depth investigation. Interestingly, all of the significantly regulated miRNAs found in this study play a role in cellular processes that have been described to be affected by lipedema, such as adipogenesis, angiogenesis, inflammation and fat metabolism.

Here, we analyzed for the first time lipedema on miRNA level. We found that sEV miRNA but not total miRNA secreted from SVF was different in healthy and lipedema patients. This study contributes to identify the role of SVF in the complex interplay of tissue components in lipedema on a miRNA level.

## Author Contributions

EP, KS, CL, MH and SW made substantial contributions to conception and design, enquired and drafted the manuscript. MB and MS as medical partner performed the patient study and collected patient data. EP, KS, MW and CL performed the scientific experiments and analyzed data. MB, JJ, JG, HR and SW have given final approval and revised the manuscript critically. All authors read and approved the final manuscript.

