## Supplemental Material for "SVF-derived extracellular vesicles carry characteristic miRNAs in lipedema"

**Suggested author order:**

Eleni Priglinger^1,2*^, Karin Strohmeier^1,2,4^, Moritz Weigl^3^, Carolin Lindner^1,2^, Martin Barsch^5^, Jaroslaw Jacak^2,4^, Heinz Redl^1,2^, Johannes Grillari^1,2,6^, Matthias Sandhofer^5^, Matthias Hackl^3#^, Susanne Wolbank^1,2#^

^#^These authors contributed equally

^1^Ludwig Boltzmann Institute for Experimental and Clinical Traumatology, AUVA Research Center, Linz/Vienna, Austria

^2^Austrian Cluster for Tissue Regeneration, Vienna, Austria

^3^TAmiRNA GmbH, Vienna, Austria

^4^School of Medical Engineering and Applied Social Science, University of Applied Sciences Upper Austria, Linz, Austria

^5^Austrian Center for Lipedema, Linz/Vienna, Austria

^6^Institute of Molecular Biotechnology, Department of Biotechnology, BOKU - University of Natural Resources and Life Sciences

**Author Email**

***Corresponding Author:**

Dr. Eleni Priglinger

Ludwig Boltzmann Institute for Experimental and Clinical Traumatology

in the AUVA trauma research center, Austrian Cluster for Tissue Regeneration.

Donaueschingenstrasse 13

A-1200 Vienna, Austria

Supplementary Figure 1

**Nanoparticle tracking analysis of conditioned medium (CM), concentrated CM (cCM) and small extracellular vesicles (sEVs).** a) Particle count of CM and cCM. b) cCM was fractionated by size exclusion chromatography. Particle count and protein measurement, obtained by absorbance at 280 nm, of obtained fractions. c) Size distribution of cCM and d) sEVs (exemplarily shown by fraction 9).

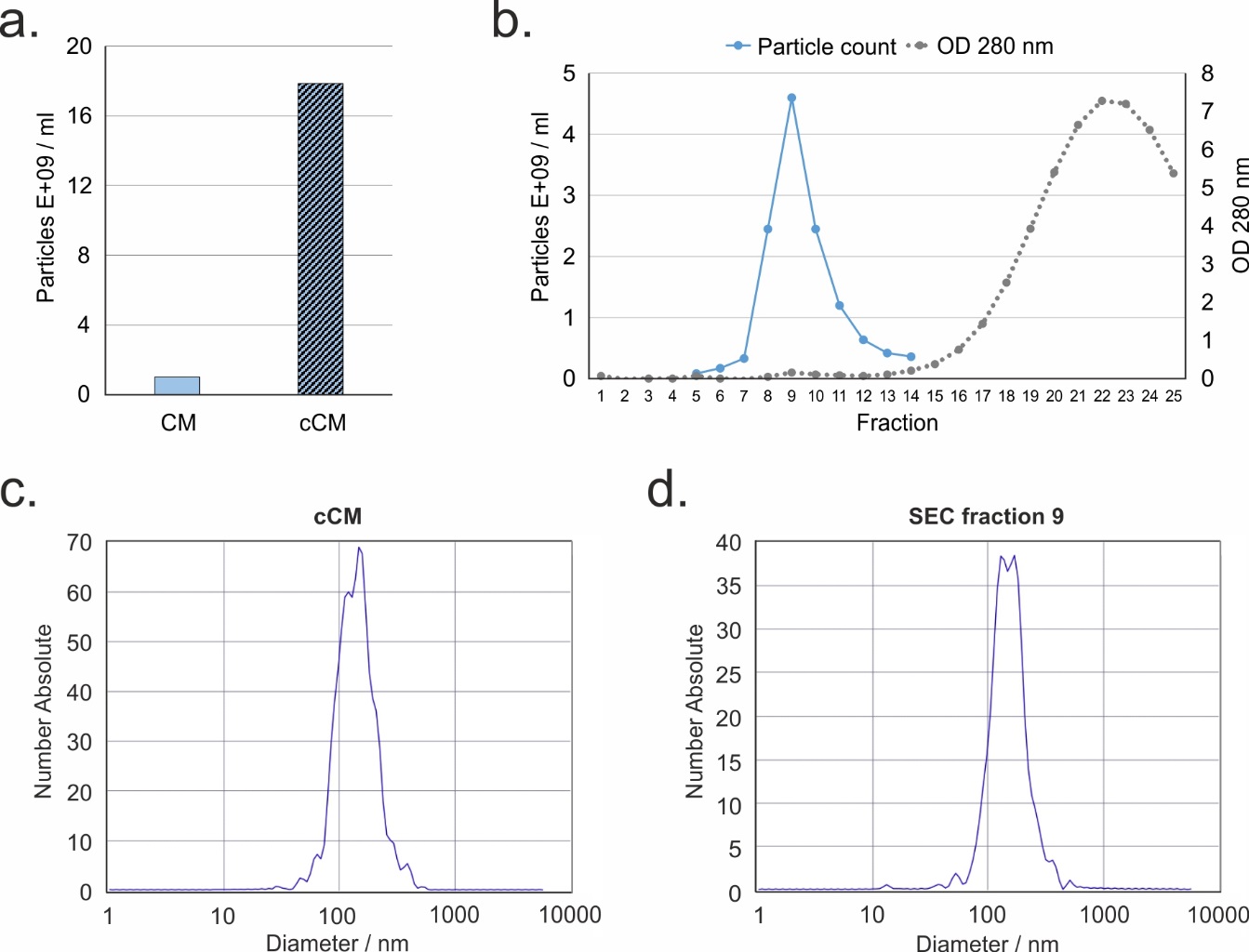

Supplementary Figure 2

**Quality control of RT-qPCR data.** Data quality for all concentrated conditioned media (cCM) and small extracellular vesicles (sEVs) samples from control (co) and lipedema (lip) was assessed using spike-in controls to determine RNA extraction efficiency, enzymatic inhibition, and overall variability. a) Cq-values for three distinct RNA spike-in controls with 100x (UniSp2), 1x (UniSp4), and 0.01x (UniSp5) concentration are shown. b) Cq-values for the spike-ins added during cDNA synthesis (cel-miR-39) and PCR amplification (UniSp3) are shown. All spike-in controls (a,b) showed homogenous values across all 12 samples with low variability.

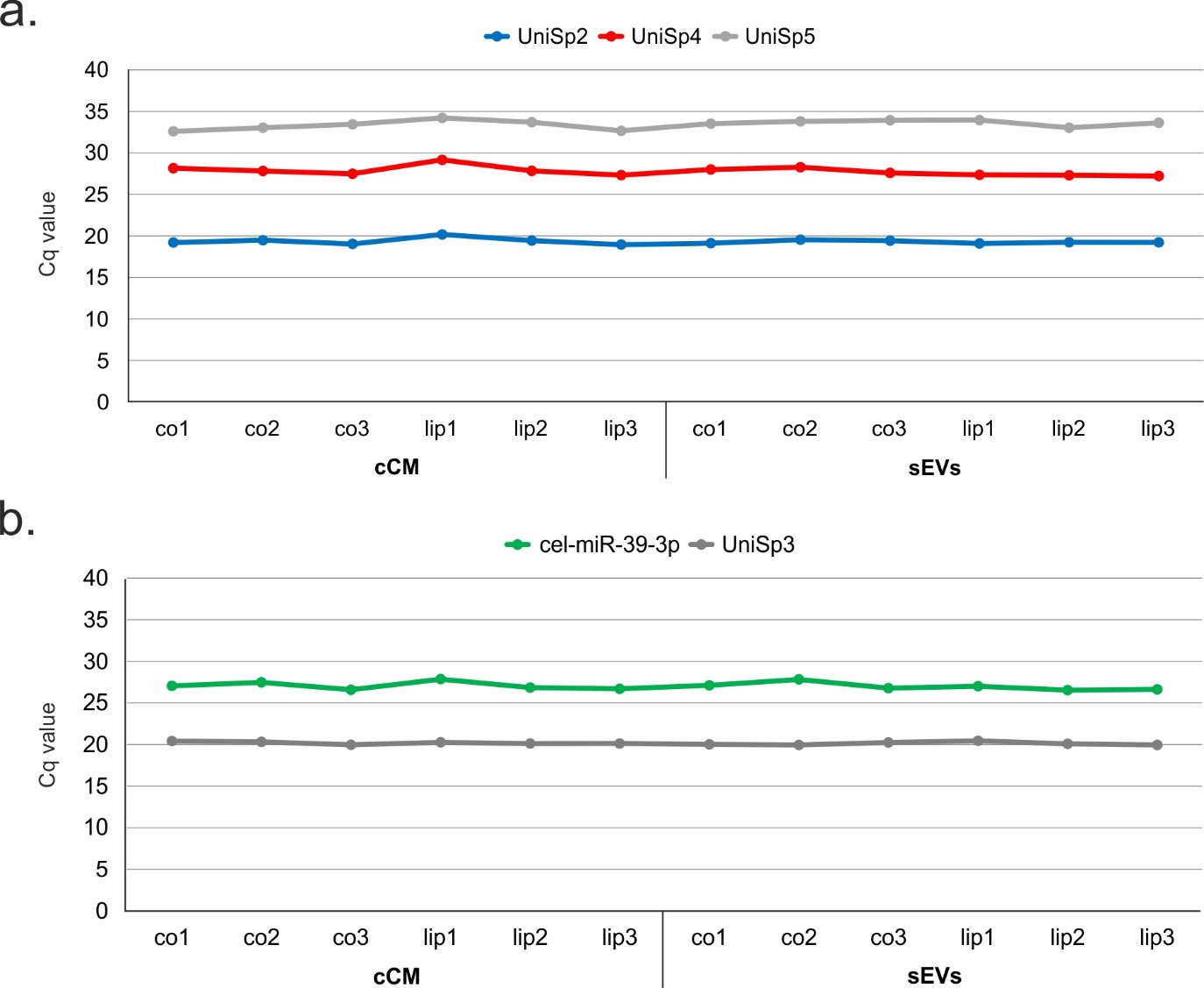

Supplementary Table 1

**Analysis of microRNA signal origin from miRNAs in small EVs.** microRNAs signals obtained from all 133 miRNAs in small extracellular vesicles (sEVs) were compared against the total microRNA signal in concentrated conditioned media (cCM) and expressed as %. For three miRNAs more than 90% of the signal in cCM originated from the sEV fraction, and for 6 further miRNAs >50% of the signal was derived from the sEV fraction. For 83 out of 133 miRNAs <10% of the total signal was obtained from the sEV fraction.

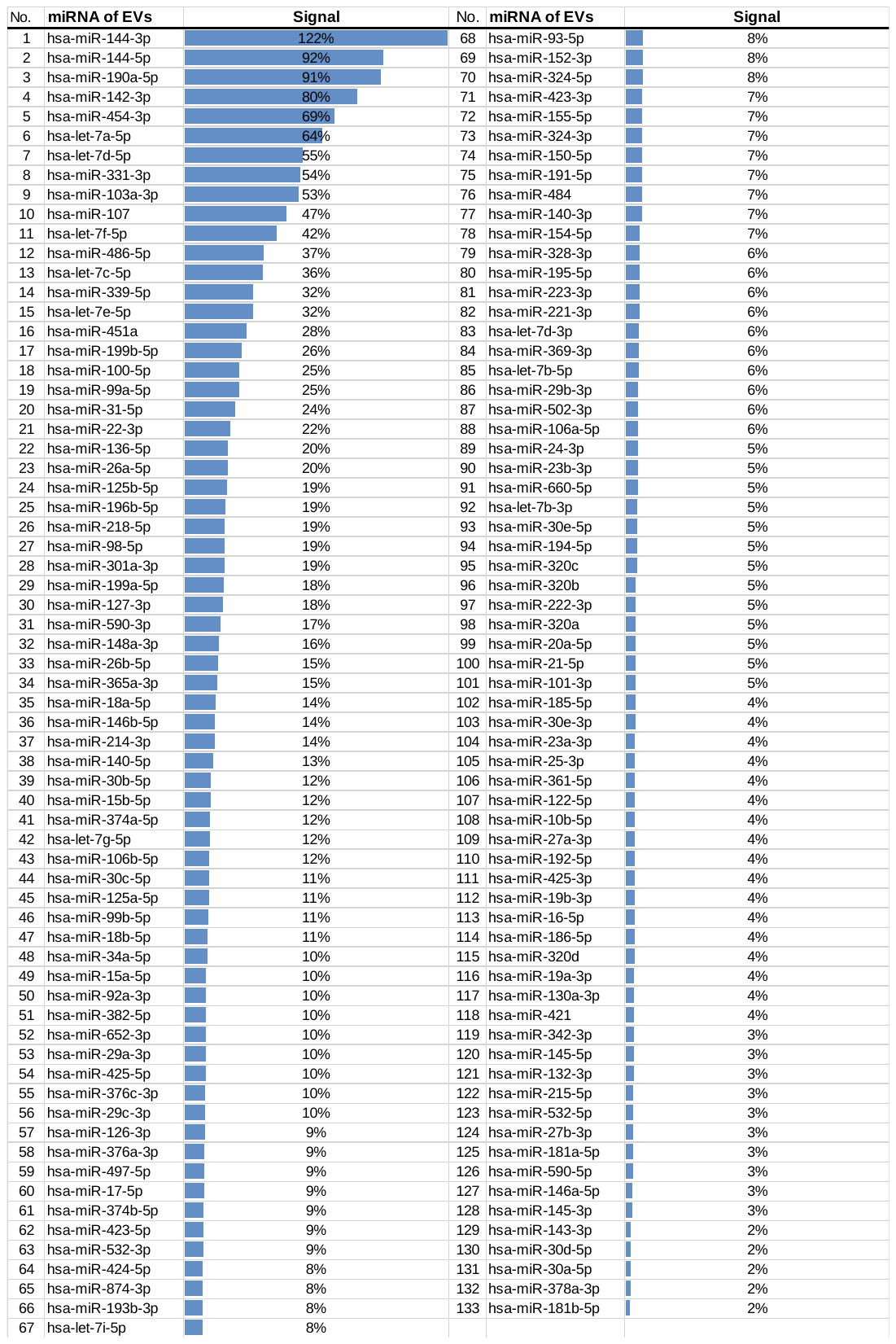
